# Miniature linear and split-belt treadmills reveal mechanisms of adaptive motor control in walking *Drosophila*

**DOI:** 10.1101/2024.02.23.581656

**Authors:** Brandon G. Pratt, Su-Yee J. Lee, Grant M. Chou, John C. Tuthill

**Affiliations:** Department of Physiology and Biophysics, University of Washington, Seattle, WA, USA

## Abstract

To navigate complex environments, walking animals must detect and overcome unexpected perturbations. One technical challenge when investigating adaptive locomotion is measuring behavioral responses to precise perturbations during naturalistic walking; another is that manipulating neural activity in sensorimotor circuits often reduces spontaneous locomotion. To overcome these obstacles, we introduce miniature treadmill systems for coercing locomotion and tracking 3D kinematics of walking *Drosophila*. By systematically comparing walking in three experimental setups, we show that flies compelled to walk on the linear treadmill have similar stepping kinematics to freely walking flies, while kinematics of tethered walking flies are subtly different. Genetically silencing mechanosensory neurons alters step kinematics of flies walking on the linear treadmill across all speeds, while inter-leg coordination remains intact. We also found that flies can maintain a forward heading on a split-belt treadmill by adapting the step distance of their middle legs. Overall, these new insights demonstrate the utility of miniature treadmills for studying insect locomotion.

## Introduction

Many animals rely on legged locomotion to move through diverse and unpredictable environments. To achieve behavioral goals in the face of this unpredictability, nervous systems have evolved to control the body in an adaptive manner. Animals as diverse as cockroaches (Couzin-Fuchs et al., 2015) and humans (Eng et al., 1994) use similar strategies to recover from unexpected motor outcomes (e.g., tripping), by rapidly adjusting coordination within and between legs. Understanding how sensorimotor neural circuits detect perturbations and generate adaptive motor responses remains a fundamental question in neuroscience (Tuthill and Wilson, 2016).

A common method to investigate the neural control of movement is to perturb neurons within candidate circuits and measure the effect on an animal’s behavior. For example, past efforts to identify sensorimotor circuits have relied on anatomical lesions (Andersson and Grillner, 1983; Dietz, 2002). While these methods revealed regions of the nervous system that are important for proprioceptive sensing and motor control, they lack cell-type specificity and produce wide-ranging behavioral effects. More recently, genetic methods have enabled targeted manipulation of specific cell-types that sense or control the body. However, these experimental manipulations often decrease the probability and vigor of spontaneous behavior. For example, the loss of feedback from mechanosensory neurons in both mammals (Chesler et al., 2016) and insects (Mendes et al., 2013) reduces walking speed and probability. This confound has made it challenging to dissect the relative roles of mechanosensory feedback vs. feedforward motor commands across different walking speeds (Bidaye et al., 2018).

One strategy to overcome this reduction in spontaneous behavior is to compel animals to walk, for example by placing them on an actuated treadmill. Treadmills have been historically used to study the neural basis of motor control and adaptive locomotion in both vertebrates (Belanger et al., 1996; Hasan and Stuart, 1988; Wetzel and Stuart, 1976) and invertebrates (Dean and Wendler, 1983; Foth and Bässler, 1985; Foth and Graham, 1983; Herreid and Full, 1984; Herreid et al., 1981; Watson and Ritzmann, 1998, 1997). For instance, treadmills have been used to drive walking in cats (Whelan, 1996) and rodents (Fujiki et al., 2018), leading to important insights into spinal circuits for adaptive locomotor control.

Because treadmills are externally controlled, they can also deliver calibrated mechanical perturbations to walking animals. Previous work showed that cats walking on a treadmill learn to increase the height of their steps to avoid being smacked by a paddle (McVea and Pearson, 2007). Split-belt treadmills, which consist of two independently controlled belts, are another classic paradigm to investigate walking coordination and motor adaptation. Both humans (Kambic et al., 2023; Reisman et al., 2007, 2005) and mice (Darmohray et al., 2019) learn new inter-leg coordination patterns when their left and right legs are driven at different speeds on a split-belt treadmill. This phenomenon of split-belt adaptation has been used to investigate behavioral and neural mechanisms of adaptive locomotion (Torres-Oviedo et al., 2011). A final advantage of treadmills is that they enable the study of locomotion within a confined space, which is important for capturing body kinematics or physiological signals from neurons and muscles.

In recent years, the fruit fly, *Drosophila melanogaster*, has emerged as an important model system for studying proprioceptive sensing and adaptive locomotion (Agrawal et al., 2020; Chen et al., 2021; Chockley et al., 2022; Isakov et al., 2016; Mamiya et al., 2023, 2018; Mendes et al., 2013). The key advantages of the fly are a compact, fully-mapped nervous system (Anthony Azevedo et al., 2022; Shin-ya Takemura et al., 2023; Sven Dorkenwald et al., 2023) and cell-type specific tools for targeted genetic manipulations. Fly locomotion has been previously studied in tethered animals walking on a floating sphere (Berendes et al., 2016; Buchner, 1976; Creamer et al., 2018; Götz and Wenking, 1973), or in freely walking animals constrained to a behavioral arena (DeAngelis et al., 2019; Fujiwara et al., 2022; Mendes et al., 2013; Simon and Dickinson, 2010; Strauss and Heisenberg, 1990; York et al., 2022). One advantage of the tethered preparation is that it enables 3D tracking of the fly’s body and legs (Günel et al., 2019; Karashchuk et al., 2021), which has not previously been possible in freely walking flies. It is also possible to record neural signals of tethered flies using optical imaging (Seelig et al., 2010) or electrophysiology (Fujiwara et al., 2017; Turner-Evans et al., 2017). However, one disadvantage of studying locomotion in tethered flies is that their posture is constrained, and normal ground reaction forces may be disrupted, which could affect walking kinematics.

To bridge these established methodologies, we introduce a new linear treadmill system that enables long-term 3D tracking of walking *Drosophila*. We systematically compare walking kinematics on the linear treadmill to those of freely walking and tethered flies. We then use the linear treadmill to investigate step kinematics and inter-leg coordination following genetic silencing of mechanosensory neurons. Last, we introduce a novel split-belt treadmill for fruit flies, which we use to uncover behavioral mechanisms of adaptive motor control. We provide open-source software and hardware designs for these treadmill systems as resources for the community.

## Results

### A linear treadmill for tracking 3D walking behavior in *Drosophila*

We engineered a miniature linear treadmill system to measure 3D walking kinematics in flies (**Figure 1A**). A fly was constrained to walk on the treadmill within a transparent chamber. Its wings were trimmed to discourage flight initiation. We used 5 high-speed video cameras (180 fps) to record fly walking behavior. To test the treadmill system, we used wild-type Berlin flies that had a body length of 2.04 ± 0.10 mm. We used DeepLabCut (Mathis et al., 2018) and Anipose (Karashchuk et al., 2021) to track and extract 3D kinematics of the fly leg tips and body (**Figure 1B**).

**Figure 1.**
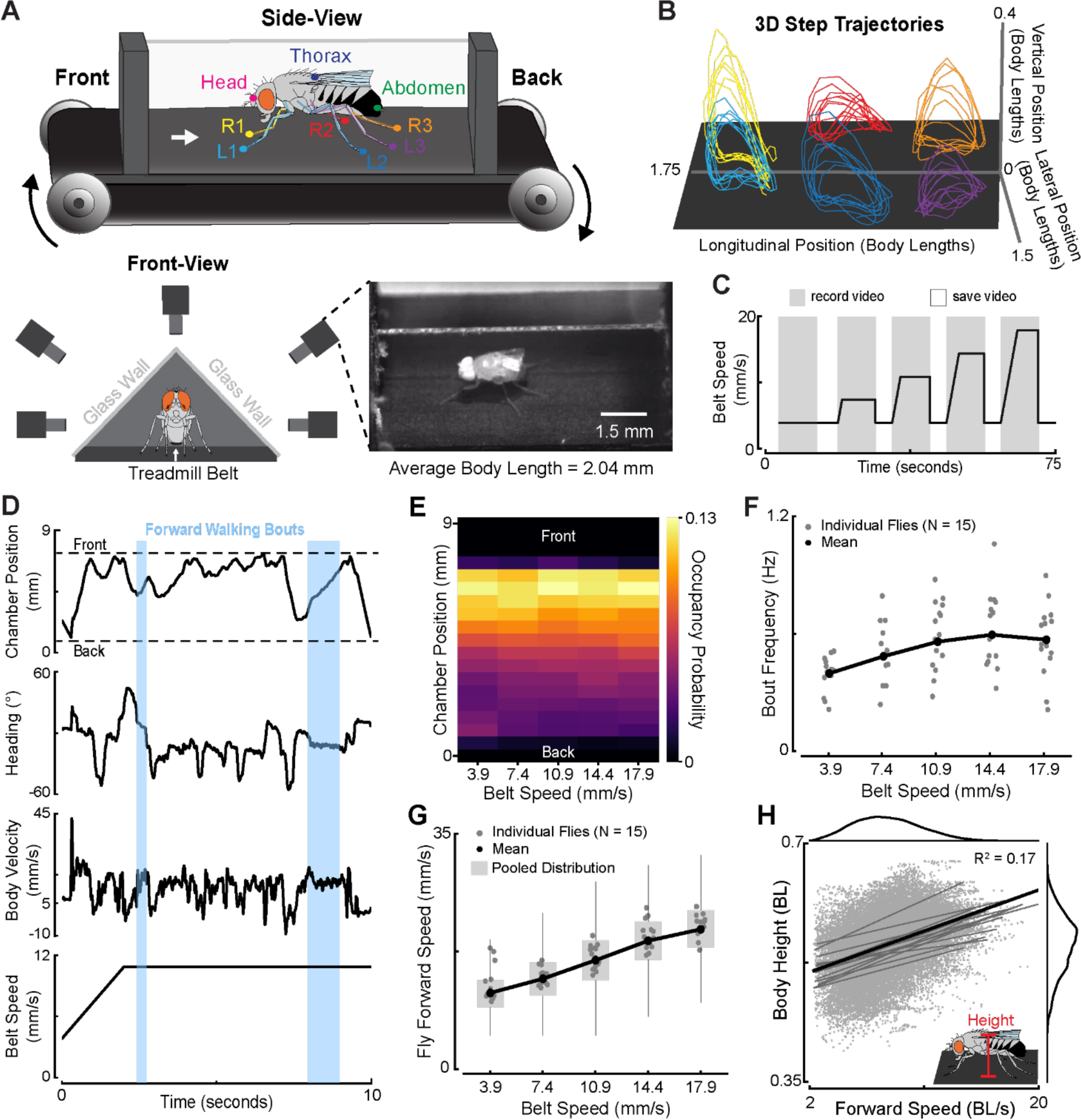
The linear treadmill controls locomotor speed and enables tracking of 3D kinematics in walking *Drosophila*. (**A**) Schematic of the linear treadmill setup. Key points associated with the head, thorax, abdomen, and each leg tip were tracked in 3D. Flies were recorded with 5 high-speed cameras as they were driven to walk on the treadmill within an attic-shaped chamber. Flies had an average body length of 2.04 ± 0.10 mm. Note that the schematic flies are not to scale. (**B**) 3D leg tip (tarsi) trajectories during forward walking. Colors correspond to the labels in (A). (**C**) Flies were compelled to walk at different speeds by moving the treadmill belt at one of 5 steady-state speeds. The belt gradually increased in speed to reach steady-state speeds greater than 5 mm/s. Each speed was presented to a fly 10 times, with speeds randomly interleaved. Each trial was composed of a 10 s recording period (gray) followed by a 5s saving period (white) in which the belt speed returned to the baseline speed. (**D**) A representative trial of a fly walking on the treadmill with a belt speed of 10.9 mm/s. The position of the fly along the chamber (measured at the thorax), the fly’s heading angle and body velocity, and the belt speed profile are plotted from top to bottom, respectively. Forward walking bouts are highlighted by the light blue shaded regions. Forward walking bouts were classified as periods lasting at least 200 ms where the fly walked in the middle of the chamber (see Methods), had a heading angle between –15 to 15 degrees with respect to the front of the chamber, and had a body velocity greater than 5 mm/s. (**E**) Flies walked between the middle and front of the chamber across all belt speeds. (**F**) Flies increased their walking bout frequency as the belt speed increased. Gray dots are the mean frequency of individual flies, while the black line denotes the mean across all flies. (**G**) Flies increased their forward walking speed as the belt speed increased. Box plots show the distribution of pooled data. Black line connects the median forward speed across all flies and belt speeds. (**H**) Flies increased their body height (vertical distance between the thorax and ground) as they walked faster (in body lengths per second, or BL/s). Black: population fit; Gray line: individual fit. See also Video 1.

We first measured the 3D kinematics of flies as they walked across a range of belt driving speeds (**Figure 1C**). As illustrated by a representative fly walking at an intermediate belt speed, flies on the treadmill displayed forward walking bouts, standing, and other lateral movements (**Figure 1D**, **Video 1**). Flies traversed the entirety of the chamber but spent most of their time toward the front (**Figure 1E**). They typically walked in short bursts, accelerating toward the front of the chamber, where they would walk for a short period, after which they would ride the belt to the back of the chamber; contact with the back of the chamber would then initiate another walking bout. Flies increased their walking bout frequency as the belt speed increased, as measured by the frequency at which they crossed the middle of the chamber from the rear (**Figure 1F**). The sporadic structure of fly treadmill walking is similar to that previously reported for freely walking flies (Sorribes et al., 2011). Flies on the treadmill also consistently increased their walking speed to keep up with the treadmill’s belt driving speed (**Figure 1G**). At the extremes, flies on the treadmill were able to sustain walking at a max belt speed of 40 mm/s (**Video 2**) and surpassed an instantaneous walking velocity of 50 mm/s (**Video 3**), which is the fastest walking speed ever recorded for *Drosophila melanogaster*. By driving flies to walk across a range of speeds while recording their 3D kinematics, we found that flies increased their body height as they walked faster by 0.007 BL per BL/s (**Figure 1H**).

In summary, our engineered treadmill makes it possible to force individual flies to walk for long periods (up to 1 hour) while tracking 3D body and leg kinematics. Flies remained upright 97% of the time on the treadmill and spent an average of 54% of the time walking, enabling collection of large amounts of useful kinematic data from each animal. Using 3D tracking, we discovered that flies elevate their body as they walked faster, a relationship that has been shown in other walking animals, from cockroaches (Full and Tu, 1991) to humans (Struzik et al., 2021), but had not been previously described in *Drosophila*.

### Comparison of walking kinematics between treadmill, freely, and tethered walking flies

Multiple previous studies have quantified step kinematics of freely walking flies (Chun et al., 2021; DeAngelis et al., 2019; Fujiwara et al., 2022; Mendes et al., 2013; Strauss and Heisenberg, 1990; Szczecinski et al., 2018; Wosnitza et al., 2013). To compare treadmill walking to freely walking kinematics, we tracked and analyzed a new dataset of wild-type flies walking in a circular arena. We focused our kinematic analyses on forward walking bouts, which were identified from fly heading direction and body velocity. Note that more detailed definitions of all kinematic parameters are included in the Methods.

We found that the key relationships between stepping kinematics and forward walking speed were similar between flies walking on the treadmill (**Figure 2i**) and freely walking flies (**Figure 2ii**, **Video 4**). Flies in both setups increased step frequency as they walked faster (**Figure 2Ai-ii**). Correspondingly, stance duration was inversely related to walking speed for flies in both setups (**Figure 2Bi-ii**). However, swing duration remained fairly constant across speeds and was of a similar magnitude for treadmill and freely walking flies (**Figure 2Ci-ii**). Step length, the distance between the footfalls of each leg, was also comparable between treadmill and freely walking flies (**Figure 2Di-ii**). Flies in both setups had similar increases in step length with increasing walking speed. The largest difference between treadmill and freely walking flies was that the step kinematics of freely walking flies were more variable (e.g., step frequency within the walking speed range of 6.5-10.1 BL/s: freely walking σ^2^ = 5.18 s^-1^; treadmill walking σ^2^ = 1.98 s^-1^). This could be because the treadmill has a single driving axis, which may result in straighter walking bouts. The step kinematics of flies in our freely walking dataset were also consistent with prior work (DeAngelis et al., 2019; Mendes et al., 2013; Strauss and Heisenberg, 1990; Szczecinski et al., 2018; Wosnitza et al., 2013), even though fly strain and sex were sometimes different in those studies (**Table S1**). Overall, the step kinematics of flies walking on the treadmill were similar to freely walking flies.

**Figure 2.**
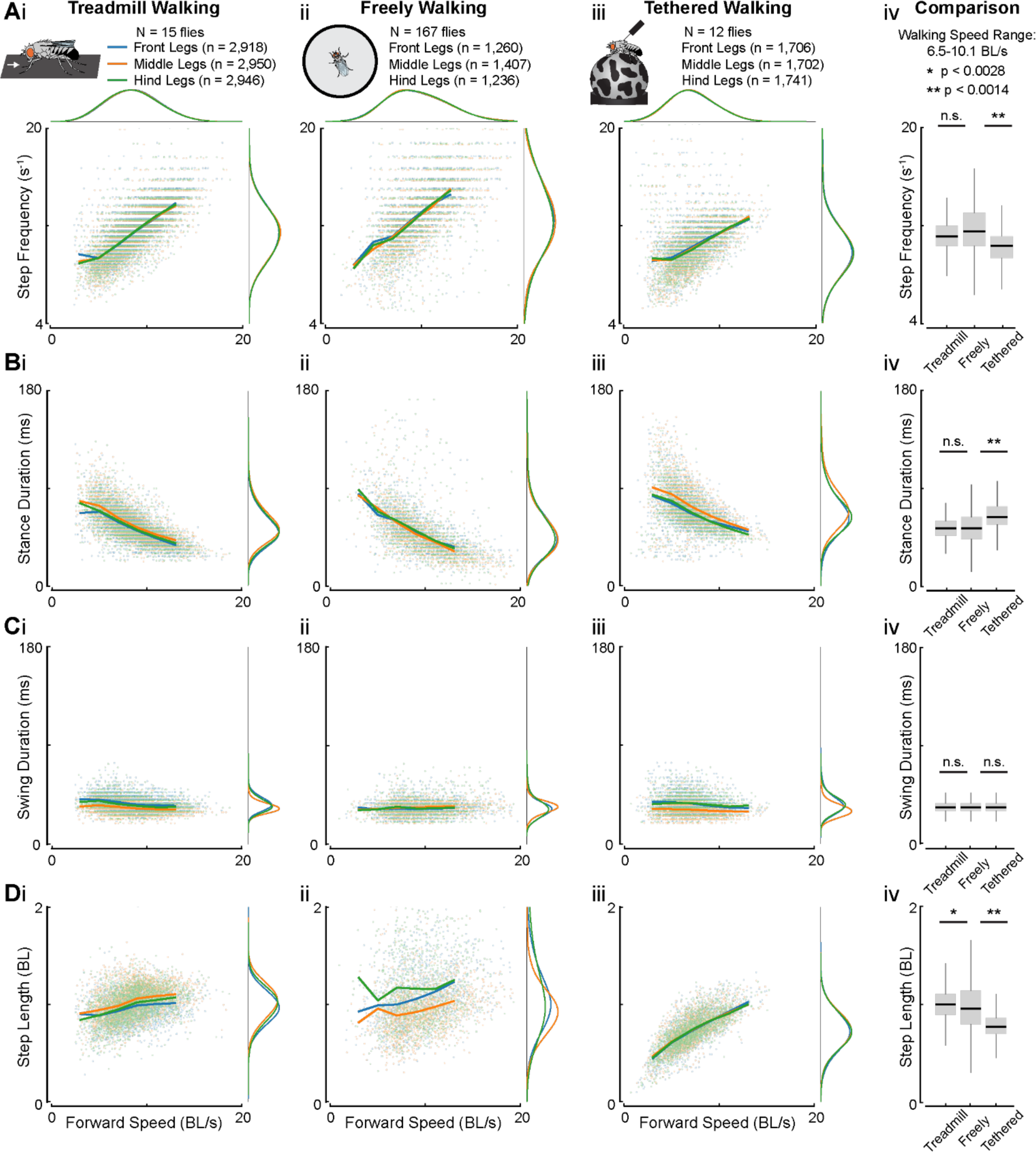
Step kinematics are similar across treadmill and freely walking flies, but different from tethered flies. (**A**) Step frequency of the front (blue), middle (orange), and hind (green) legs as a function of forward walking speed for treadmill (i), freely walking (ii), and tethered flies (iii). Distributions (iv; gray box plots) that combine step frequency across leg pairs for treadmill, freely, and tethered walking flies over an overlapping and dense range of walking speeds (6.5-10.1 BL/s). (**B**) Stance duration as a function of forward walking speed for flies in the different setups. (**C**) Swing duration as a function of forward walking speed. (**D**) Step length as a function of forward walking speed. In **A-D,** lines are speed-binned averages (2 BL/s bins between 3 and 13 BL/s) for each kinematic parameter and leg pair. Step frequency, stance duration, and swing duration (**A-C**) were computed on interpolated data for treadmill (i; 180 to 300 fps) and freely walking flies (ii; 150 to 300 fps) to enable the comparison to tethered flies. Distributions associated with each leg were visually offset from each other for presentation in **A-C**. In **A-C iv**, Chi-squared test for goodness of fit with a Bonferroni correction of 18 was used to statistically compare the distributions across the different walking setups. In **D**, t-test with a Bonferroni correction of 18 was used to compare the distributions. N: number of flies; n: number of steps. See also Figure S1 for comparisons of the similarity between the setups with respect to step kinematics, leg pairs, and walking speeds.

Having developed a framework for comparing walking kinematics across experimental setups, we took the opportunity to extend our analysis to a third setup: tethered flies walking on a floating sphere (**Figure 2iii**, **Video 5**). The “fly-on-a-ball” setup is commonly used in our lab (Agrawal et al., 2020; Azevedo et al., 2020; Karashchuk et al., 2021) and many others (e.g., Berendes et al., 2016; Creamer et al., 2018; Seelig et al., 2010), but leg kinematics of tethered and freely walking flies have not been systematically compared.

In general, we found that the relationships between stepping kinematics and walking speed were similar across all three setups. However, tethered flies differed from untethered ones (i.e. treadmill and freely walking flies) in several key aspects. First, tethered flies didn’t reach the faster walking speeds displayed by untethered flies. Step frequency of tethered flies was also significantly lower than that of untethered flies (**Figure 2Aiv**), whereas stance duration was significantly longer (**Figure 2Biv**). There was no significant difference in swing duration across all setups (**Figure 2Civ)**. Therefore, the reduced step frequency in tethered flies resulted from longer stance durations across walking speeds. Tethered flies also had a significantly lower step length than untethered flies (**Figure 2Div**). However, the step length of tethered flies was more correlated with walking speed (tethered: r = 0.41; freely: r = 0.05; treadmill: r = 0.22). We also compared step kinematics between the different setups across leg pairs and walking speeds (**Table S1**). We found that there was a greater similarity in the step kinematics between treadmill and freely walking flies, especially at fast walking speeds. These differences in step kinematics and walking speed ranges between tethered and untethered flies may be because the tethered flies are walking on a spherical surface and/or because their body weight is supported by a rigid tether.

We next compared inter-leg coordination across the three setups. We quantified the number of legs that were in the stance phase of the step cycle at each point in time, which remains constant for idealized coordination patterns, such as the tripod coordination pattern where 3 legs are in stance (DeAngelis et al., 2019). We found that the probability of having a specific number of legs in stance was similar between tethered and untethered flies. Specifically, flies in all three setups showed an increased probability of having 3 legs in stance as walking speed increased (**Figure 3A**). This is consistent with prior work showing that freely walking flies are more likely to use a canonical tripod coordination pattern at higher speeds (DeAngelis et al., 2019; Mendes et al., 2013; Pereira et al., 2019; Wosnitza et al., 2013).

**Figure 3.**
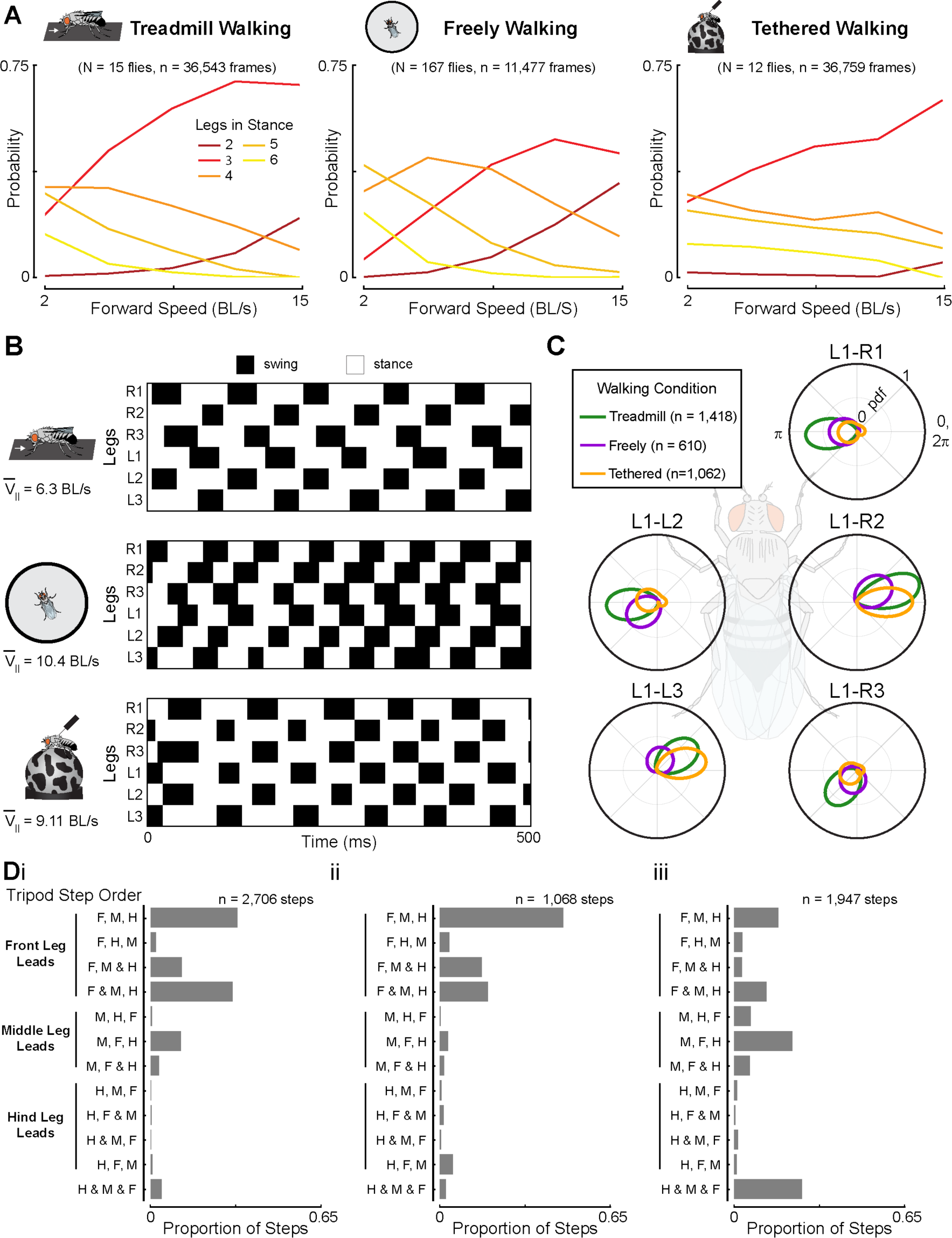
Subtle differences in inter-leg coordination between tethered and untethered flies. (**A**) Probability of treadmill, freely walking, and tethered flies exhibiting 2-6 legs in stance across forward walking speeds. Lines are speed-binned averages of the probability of a certain number of legs in stance (3.2 BL/s bins between 2 and 15 BL/s). N: number of flies; n: number of camera frames. (**B**) Representative plots of the swing and stance phases of each leg across all setups. Black: swing; White; stance. **(C)** Polar plots of the relative phase between the left front leg and each other leg for flies in each setup. Kernel density estimations were used to determine the probability density functions. n: number of phase comparisons. (**D**) Bar plot of the proportion of steps attributed to each step order combination of the legs with a tripod for treadmill (i), freely (ii), and tethered (iii) walking flies. The leading leg is indicated on the left of the graph. ‘&’ denotes legs contacting the ground at the same time. n: number of steps.

Given that flies across all three setups had similar speed-dependent changes in inter-leg coordination, we next asked if the stepping pattern underlying inter-leg coordination was different between tethered and untethered flies. We therefore examined the relative swing and stance relationships across legs. We observed that the order in which legs entered stance conformed to the so-called *Cruse Control* rules for both tethered and untethered flies (Bidaye et al., 2018; Cruse, 1990, 1985). This is illustrated by the anterior progression of ipsilateral leg stepping (diagonal black stripes in **Figure 3B**). We also examined the relative phase relationships between the left front leg and all other legs across the different setups. we computed phase by determining when a leg entered stance within the left front leg’s step cycle. We found that the relative phase relationships across all legs were similar for treadmill, freely, and tethered walking flies (**Figure 3C**). However, the phase relationships of the ipsilateral front and hind legs and contralateral middle leg that make up the canonical tripod were more coupled for tethered flies.

Finally, we looked at the order in which legs within a tripod entered stance with respect to the left front leg’s step cycle. We found differences in the stance onset order between tethered and untethered flies. For example, the front leg within a tripod was usually the first leg to contact the ground for treadmill (**Figure 3Di**) and freely walking flies (**Figure 3Dii**), whereas the stance order was more variable for tethered flies, having more instances of the middle leg entering stance first (**Figure 3Diii**). Tethered flies also had more instances where all legs within a tripod entered stance at the same time, called an “ideal tripod”. Therefore, the inter-leg phase coupling of tethered flies was stronger than that of untethered flies. One explanation could be that the added stability from the tether induces a more tightly coupled inter-leg coordination pattern.

In summary, we found that although step kinematics were subtly different between tethered and untethered flies (**Figure 2**), inter-leg coordination was broadly similar (**Figure 3**).

### Silencing mechanosensory feedback alters step kinematics across walking speeds but not inter-leg coordination

One of our motivations to develop a linear treadmill was to investigate the role of mechanosensory feedback in fly locomotion. This has historically been challenging, because silencing mechanosensory neurons typically leads to a reduction in locomotor probability and speed in flies (Mendes et al., 2013) and other animals (Chesler et al., 2016; Dietz, 2002). To test whether flies lacking mechanosensory feedback will walk on the linear treadmill, we genetically silenced chordotonal neurons (iav-GAL4 > UAS-kir2.1), which are found at multiple joints throughout the fly’s body, including in the femur and tibia (**Figure 4A**). As expected, silencing chordotonal neurons drastically reduced locomotion in freely walking flies (**Figure 4B**). However, flies lacking chordotonal feedback walked a greater proportion of the time and across a wider range of speeds when driven to walk on the linear treadmill (**Figure 4C**). Silencing chordotonal neurons altered the structure of fly locomotion on the treadmill compared to genetically-matched controls. In particular, flies with silenced chordotonal neurons spent more time towards the back of the chamber at faster belt speeds (**Figure 4D**) and exhibited a lower bout frequency (**Figure 4E**).

**Figure 4.**
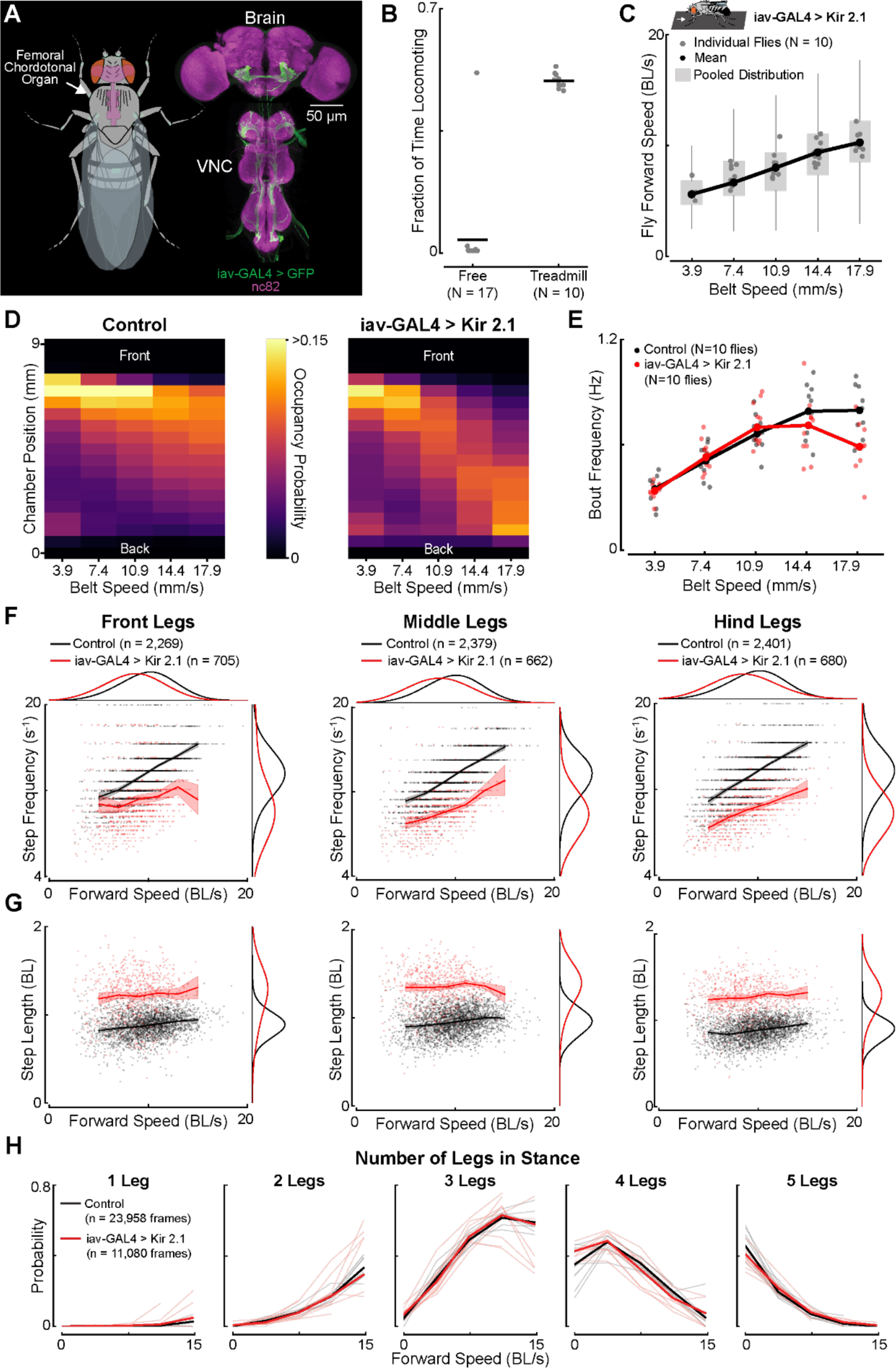
**Silencing mechanosensory chordotonal neurons alters step kinematics but not inter-leg coordination across walking speeds**. (**A**) A schematic of the locations of chordotonal neurons (green), including the femoral chordotonal organ in each leg (white arrow for left front leg), labeled by the iav-GAL4 driver line. The axons of chordotonal neurons (green) are shown in the max intensity projection of a confocal stack of the fly brain and VNC (magenta). (**B**) Flies with silenced chordotonal neurons (iav-GAL4> Kir 2.1) spent more walking on the treadmill compared to the arena. Gray dots: individual flies; Black lines are means. (**C**) The flies in (B) increased their forward walking speed as the belt speed increased. N: number of flies. Box plots show the distribution of pooled data. Black line connects the median forward speed for all flies for each belt speed. Gray dots are the median forward speed of each fly. (**D**) Heatmap of the occupancy probability along the chamber of genetically-matched controls (R52A01 GAL4-DBD > Kir2.1; left panel) and flies lacking chordotonal feedback (right) shows that the latter has a greater probability of being located towards the back of the chamber at fast belt speeds. (**E**) Bout frequency was lower at fast belt speeds for flies lacking chordotonal feedback (red) compared to controls (black). Dots are the mean bout frequency for each fly. N: number of flies. (**F**) The step frequency of the front (left), middle (center), and hind (right) legs was lower across walking speeds for flies lacking chordotonal feedback (red) compared to genetically-matched controls (black). Lines are speed binned averages (2 BL/s bins from 5-15 BL/s) for each kinematic parameter and the 95% confidence interval is shown by the shaded region. Distributions between control and experimental flies were offset from each other so that they could be visualized. n: number of steps. (**G**) Step length was greater across walking speeds for flies lacking chordotonal feedback (red) compared to controls (black). (**H**) The probability of a certain number of legs in stance (1-5: left-right) was not different across walking speeds between flies with silenced chordotonal neurons (red) and controls (black). Solid lines: pooled data; Thin lines: individual flies. n: number of camera frames.

The leg movements of flies with silenced chordotonal neurons were noticeably different. Qualitatively, the legs appeared less rigid and moved with less precision (**Video 6**). Therefore, we next analyzed the impact of silencing chordotonal neurons on step kinematics and inter-leg coordination. We found that flies lacking chordotonal feedback had a lower step frequency across legs and speeds compared to control flies (**Figure 4F**). In addition, flies with silenced chordotonal neurons had greater step lengths across speeds, suggesting that they increased the size of their steps to compensate for taking fewer of them (**Figure 4G**). However, inter-leg coordination was not significantly different between controls and flies lacking chordotonal feedback (**Figure 4H**).

In summary, we found that silencing mechanosensory feedback from chordotonal neurons altered step kinematics across all walking speeds but did not have a significant effect on inter-leg coordination. We conjecture that other proprioceptor classes, such as hair plates and campaniform sensilla (Tuthill and Azim, 2018), may play a more important role than chordotonal neurons in inter-leg coordination. Overall, these results also demonstrate the advantage of the linear treadmill to investigate the role of sensory feedback in locomotion.

### A split-belt treadmill reveals that middle legs correct for rotational perturbations

We next engineered a split-belt treadmill to investigate behavioral mechanisms of adaptive motor control in walking flies (**Figure 5A**). We compared the leg kinematics of walking flies while the belts moved at the same speed (tied) vs. when the belts moved at different speeds (split: slow and fast), focusing our analysis on periods of forward, straight walking (see Methods, **Figure 5B left**, **Video 7**). We tested splits in both directions and pooled symmetric conditions for subsequent analyses.

**Figure 5.**
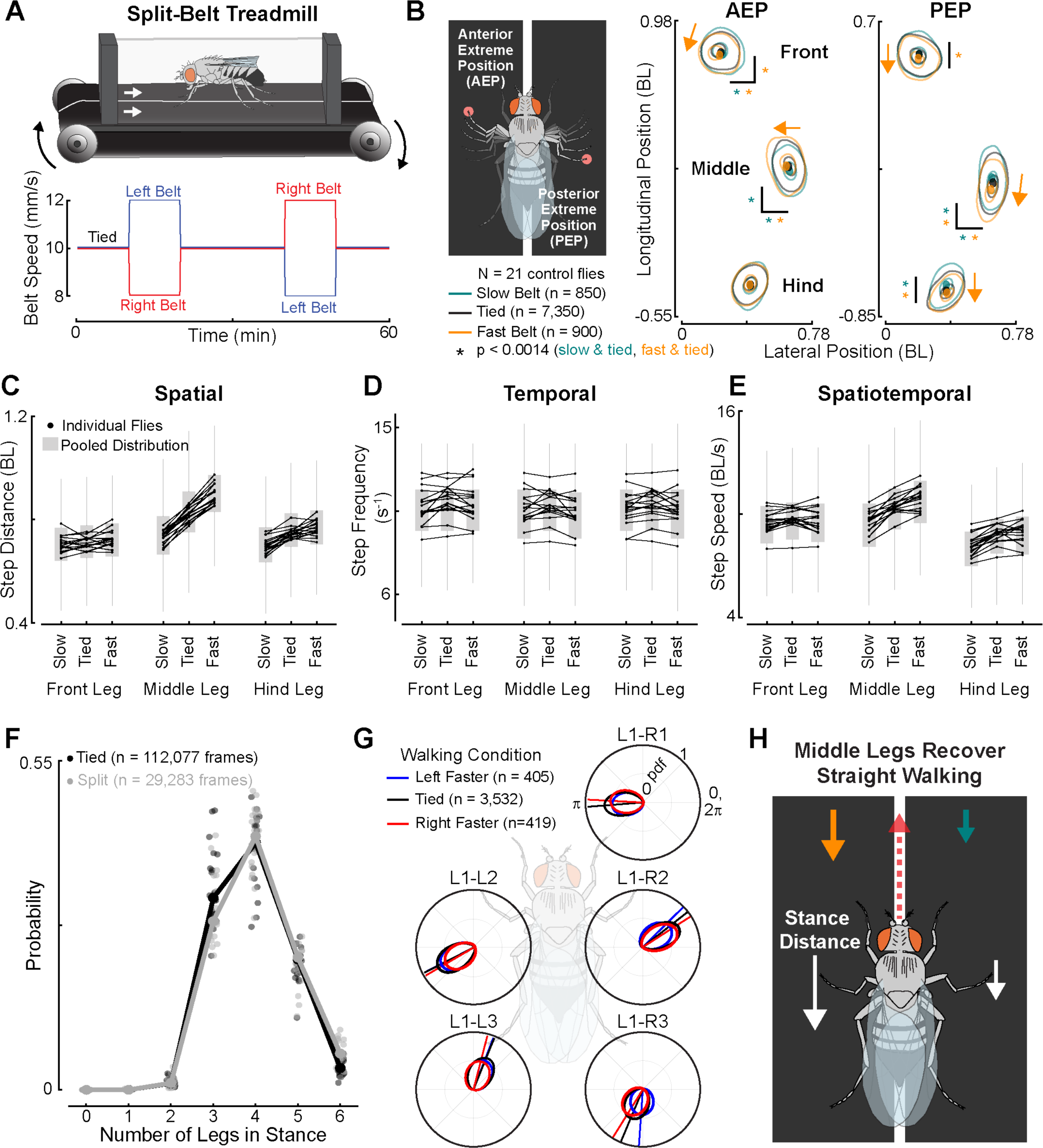
A split-belt treadmill reveals that flies adjust spatial kinematics to correct for rotational perturbations. (**A**) A schematic of the split-belt treadmill (top) and the belt speed protocol (bottom). The split-belt treadmill consists of two independently controlled belts, which were initially driven at the same speed (tied). The right (red) and left (blue) belts then differed in speed by 40%, and the direction of the speed change reversed on the subsequent split period. (**B**) We focused our analysis on forward walking bouts where the left and right legs moved on their respective belts (left). The distributions of where the legs first contact the ground, the anterior extreme positions (AEP), and where they takeoff from the ground, the posterior extreme positions (PEP), are shown for when legs walked on belts during the tied condition (black) or on the slow (teal) and fast (orange) belt during the split condition. Distributions are shown by kernel density estimations and means are denoted by dots. Arrows indicate the direction the distributions shifted with respect to the tied distribution. Bootstrapping with a Bonferroni correction of 36 was used to statistically compare the means of the tied and split distributions. An absence of a “*” indicates no significant difference. N: number of flies; n = number of steps. (**C**) Step distance increased for the middle leg as belt speed increased. The hind leg had a lower step distance when walking on the slow belt. Box plots show the distribution of pooled data. Black lines and dots are the median step distances of individual flies. (**D**) Step frequency did not change across conditions for any leg. (**E**) Step speed increased for the middle leg as belt speed increased and was lower for the hind leg on the slow belt. (**F**) Probability of 0-6 legs being stance when the belts were tied (black) or split (gray) in speed. Dots connected by lines are the global probabilities. Individual dots are the probability that a given fly shows a certain number of legs in stance. (**G**) Polar plots of the relative phase between the left front leg and the other legs when the belts were tied (black) or split (left faster: blue; right faster: red) in speed. n: number of phase comparisons. (**H**) Summary schematic showing that the middle legs adjust their step distance to recover straight walking in the presence of asymmetric belt speeds.

We found that flies achieved straight walking by modifying spatial rather than temporal step kinematics. For instance, flies significantly altered where the legs contacted or lifted off of the ground (i.e., the anterior and posterior extreme positions (**Figure 5B**). Specifically, the mean anterior extreme position (AEP) shifted posteriorly and medially for the front leg on the fast belt, and laterally for the front leg on the slow belt. The front leg’s mean posterior extreme position (PEP) also shifted posteriorly when on the fast belt. The middle leg’s AEP shifted medially when on the fast belt but translated posteriorly and laterally on the slow belt. Meanwhile, the PEP of the middle legs shifted in the opposite direction. The hind leg shifted its mean PEP in a similar manner along the longitudinal axis. These changes in leg placement produced changes in the total distance that the legs, particularly the middle legs, traveled during a step cycle with respect to the body (i.e., step distance, **Figure 5C**). Step frequency, the timing component of the step cycle, did not change (**Figure 5D**), which suggests that changes in step distance alone dictated the speed at which legs moved through their step cycle (**Figure 5E**). For example, the increased step distance of the middle leg on the fast belt resulted in a faster step speed. Although changes in step speed could in principle have been achieved by altering temporal kinematics, spatial kinematics, or a combination of both, our results suggest that flies overcome the belt asymmetries to maintain forward locomotion by specifically adjusting the step size of the middle legs.

Split-belt walking had minimal effects on inter-leg coordination. For instance, there was only a small difference in the probability of 3 legs being in stance during asymmetric belt movement compared to when the belts were tied in speed (**Figure 5F**). Moreover, the mean phase offsets between legs were either not altered or shifted slightly when the belts were driven at different speeds compared to when they were tied (**Figure 5G**). These subtle changes in inter-leg coordination may help correct for the rotational perturbation induced by the treadmill. However, the more substantial changes in the spatial positioning of the middle legs appear to be the primary mechanism that flies use to achieve straight walking during split-belt walking (**Figure 5H**).

## Discussion

In this study, we engineered miniature linear and split-belt treadmills for walking *Drosophila*. Flies walking on the treadmill exhibited similar walking behavior, step kinematics, and inter-leg coordination to freely walking flies. The treadmill allowed us to achieve the first 3D tracking of untethered fly walking, which revealed that flies elevate their body height as they walk faster. We also used the linear treadmill to show that flies lacking mechanosensory feedback from chordotonal neurons are able to walk at higher speeds if compelled to do so. Across all walking speeds, silencing chordotonal neurons altered motor control of individual legs, but not coordination between legs. Finally, we found that flies can maintain a forward heading on a split-belt treadmill by adjusting the step size of their middle legs. These insights illustrate how treadmills fill an important gap between freely walking and tethered preparations for investigating neural and behavioral mechanisms of fly locomotion.

Although our primary goal was to compare treadmill and freely walking flies, we also took the opportunity to examine walking kinematics of tethered flies. The advantage of tethered walking is that it enables full 3D joint tracking (Günel et al., 2019; Karashchuk et al., 2021), spatially targeted optogenetic stimulation (Agrawal et al., 2020), and recordings of neural activity with calcium imaging (Seelig et al., 2010) or electrophysiology (Fujiwara et al., 2017; Turner-Evans et al., 2017). However, our results suggest that studies of tethered walking should be interpreted with caution, because walking kinematics of tethered flies differ in subtle but important ways from untethered flies (**Figures 2-3**; **Figure S1**). Although speed-dependent changes in walking kinematics and coordination were consistent between tethered and untethered flies, the magnitude of step kinematics and the coupling strength between legs were different. One reason for these differences may be that tethered flies walk on a sphere (i.e., a foam ball), whereas treadmill and freely walking flies walk on a flat surface. Tethered flies also do not support their own body weight, but instead use their legs to rotate the floating sphere, a configuration that is unlikely to mimic normal ground reaction forces. Indeed, prior work showed that changing the load on the body alters walking kinematics in freely walking flies (Mendes et al., 2014). On the other hand, it is remarkable that flies and other animals walk at all while tethered on a floating sphere. Given the major mechanical differences between tethered and untethered conditions, the kinematic differences we found are relatively subtle.

One of our primary motivations for developing a treadmill system for flies was to investigate the role of mechanosensory feedback across walking speeds. Prior work has suggested that proprioception is most important at slower walking speeds, and that flies use a more feedforward motor program when walking faster (Bidaye et al., 2018). However, testing this hypothesis has been challenging, because silencing mechanosensory neurons causes flies to walk less and at lower velocities (Mendes et al., 2013). The linear treadmill makes it possible to drive fly locomotion across a wide range of speeds, including after genetic manipulations to mechanosensory neurons. We found that silencing mechanosensory feedback alters step kinematics across all walking speeds. Indeed, the greatest deviation in step kinematics between control and experimental flies occurred at faster walking speeds. However, the lack of mechanosensory feedback did not impact inter-leg coordination at any walking speed. We chose to use a blunt manipulation with a broad driver line (iav-Gal4) to illustrate the utility of the treadmill for driving walking even when mechanosensory feedback is profoundly altered. One important caveat is that we expressed Kir 2.1 in chordotonal neurons throughout development, so the nervous system could have compensated to maintain normal inter-leg coordination. In the future, it will be interesting to silence mechanosensory feedback of flies walking on the treadmill with more temporal and genetic specificity, for example using optogenetic manipulation of proprioceptor subtypes in the femoral chordotonal organ (Chen et al., 2021; Chockley et al., 2022). Our results are consistent with the hypothesis that proprioceptors in the femoral chordotonal organ contribute to individual leg kinematics, whereas descending commands from the brain (Bidaye et al., 2018) or feedback signals from other proprioceptor classes may have more influence on inter-leg coordination (Tuthill and Azim, 2018).

In hexapod insects, each pair of legs plays a specialized role in controlling locomotion. In freely walking insects, the front legs are typically used for steering (Isakov et al., 2016), while the hind legs contribute to propulsion and jumping (Burrows, 2013, 2007; Card and Dickinson, 2008). Using the split-belt treadmill, we found that the middle legs play a unique role in correcting for perturbations that displace flies from a forward walking trajectory. The middle legs are ideally positioned to stably pivot the body of the fly about its center of mass, like rowing a boat from its center. In larger insects, the middle legs have been shown to play a role in executing tight turns (Cruse et al., 2009). Although the split-belt treadmill puts the fly in artificial circumstances, it mimics many situations in the wild when flies may need to perform rotational body corrections. For example, heterogeneous terrain, meddlesome conspecifics, or unilateral wind gusts could asymmetrically act on the movement of the left and right legs to induce a rotation of the body. In the future, the split-belt treadmill may also provide a useful method to study the neural mechanisms that underlie adaptive heading stabilization in walking flies (Haberkern et al., 2022).

In addition to their utility for investigating sensorimotor control of fly walking, we anticipate several additional applications of miniature treadmill systems. One will be to investigate motor adaptation during split-belt walking, a phenomenon which has been extensively studied in mammals (Hinton et al., 2020). The split-belt treadmill is also used as a clinical tool for diagnosing cerebellar deficits (Hoogkamer et al., 2015) and post-stroke rehabilitation (Reisman et al., 2007) in humans. The linear treadmill may also be useful for the study of insect respiratory physiology, which has previously been studied during flight (Lehmann, 2001) and in running cockroaches (Herreid and Full, 1984). Finally, because our treadmill system is constructed of simple and inexpensive belts, pulleys, and motors, it can be easily customized to study other walking insects, such as ants (Dahmen et al., 2017) and snow flies (Golding et al., 2023).

## Acknowledgements

We thank Max Mauer for engineering the first prototype of the treadmill, Sarah Walling-Bell and Srinidhi Naidu for performing some of the initial treadmill experiments, Eric Martinson for valuable advice and technical support throughout the development of the treadmill systems, Tom Daniel and Jeff Riffell for letting us use their 3D printing equipment and providing advice on the development of the treadmill chamber, Michael Dickinson for designing the fly cartoons many years ago and reminding us of it often, Anne Sustar for the brain and VNC images in Figure 4A, and members of the Tuthill and B.W. Brunton labs for feedback on the manuscript. B.G.P was supported by an NSF Graduate Research Fellowship (Fellow ID: 2018261272). S.J.L. was supported by T32 NS 99578-3. Other support was provided by National Institutes of Health grants R01NS102333 and U19NS104655, a Searle Scholar Award, a Klingenstein-Simons Fellowship, a Pew Biomedical Scholar Award, a McKnight Scholar Award, a Sloan Research Fellowship, the New York Stem Cell Foundation, and a UW Innovation Award to J.C.T. J.C.T is a New York Stem Cell Foundation – Robertson Investigator.

## Author Contributions

B.G.P. and J.C.T. conceived of the study. B.G.P. engineered the linear and split-belt treadmills. B.G.P. developed high-speed videography acquisition scripts and code to control the treadmill belts. B.G.P. collected and analyzed linear and split-belt treadmill data. S.J.L. collected and analyzed freely walking data. G.M.C. collected and analyzed tethered walking data. B.P. and G.M.C. performed the statistical analyses. B.G.P. and J.C.T. wrote the manuscript with input from other authors.

## Data and Code Availability

Data is available on Dryad (https://doi.org/10.5061/dryad.mpg4f4r73). Code for analyzing and visualizing treadmill, freely, and tethered walking kinematics is located on GitHub (https://github.com/Prattbuw/Treadmill_Paper). The. stl file used for the 3D printed treadmill chamber and Python scripts to acquire high-speed video and control the treadmills are also located there.

## Supplemental information

**Figure S1.**
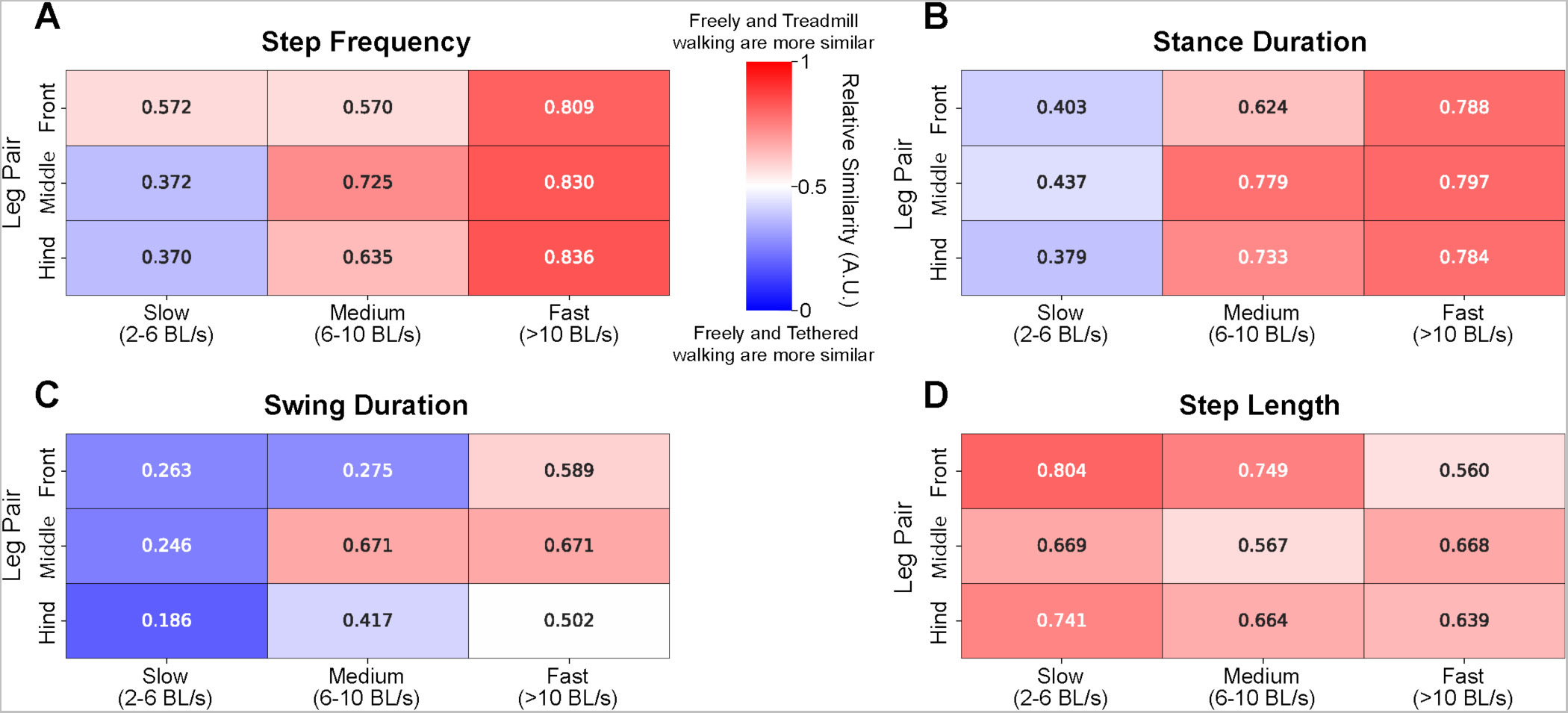
**Distributions of step kinematics are more similar between freely and treadmill walking flies than between freely walking and tethered flies**. (**A**) Step frequency was more similar between treadmill and freely walking flies, especially at medium (6-10 BL/s) and fast (>10 BL/s) walking speeds. At slow (2-6 BL/s) walking speeds, freely and tethered walking step frequencies were more similar. Relative similarity was determined by dividing the KL divergence of freely and treadmill walking kinematic distributions by the sum of the KL divergences between freely and treadmill walking, and freely and tethered walking kinematic distributions. We then reversed the similarity scale by computing 1 minus these values. A value close to 1 indicates that freely and treadmill step kinematics are more similar than freely and tethered walking kinematics for a given walking speed range. The opposite is true for values close to 0. (**B**) Stance duration was more similar between freely and treadmill walking. At slow walking speeds, freely and tethered walking stance durations were slightly more similar than that of treadmill walking. (**C**) Swing duration was more similar between freely and treadmill walking flies at fast speeds, whereas the swing duration of freely and tethered walking flies was more similar at slower speeds. (**D**) Step length was most similar between freely and treadmill walking flies across all speeds and legs.

**Table S1.**
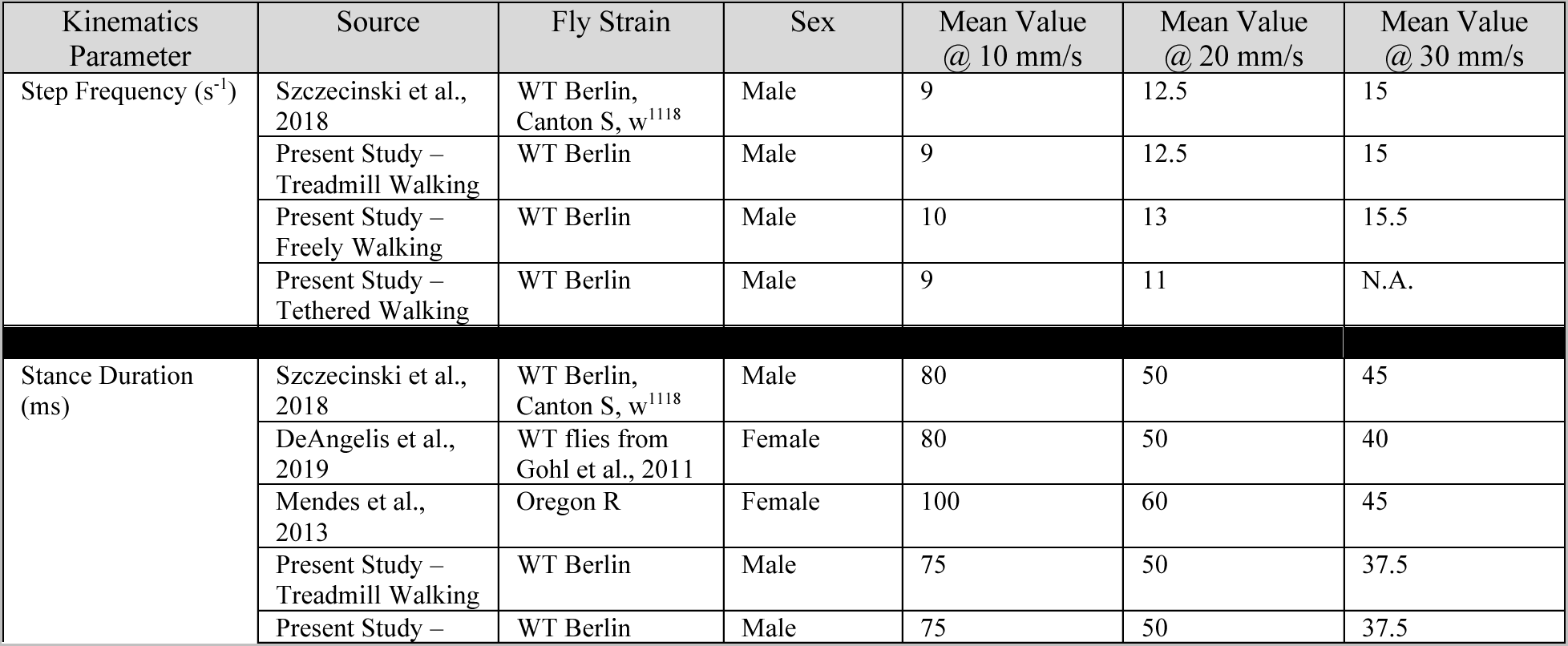
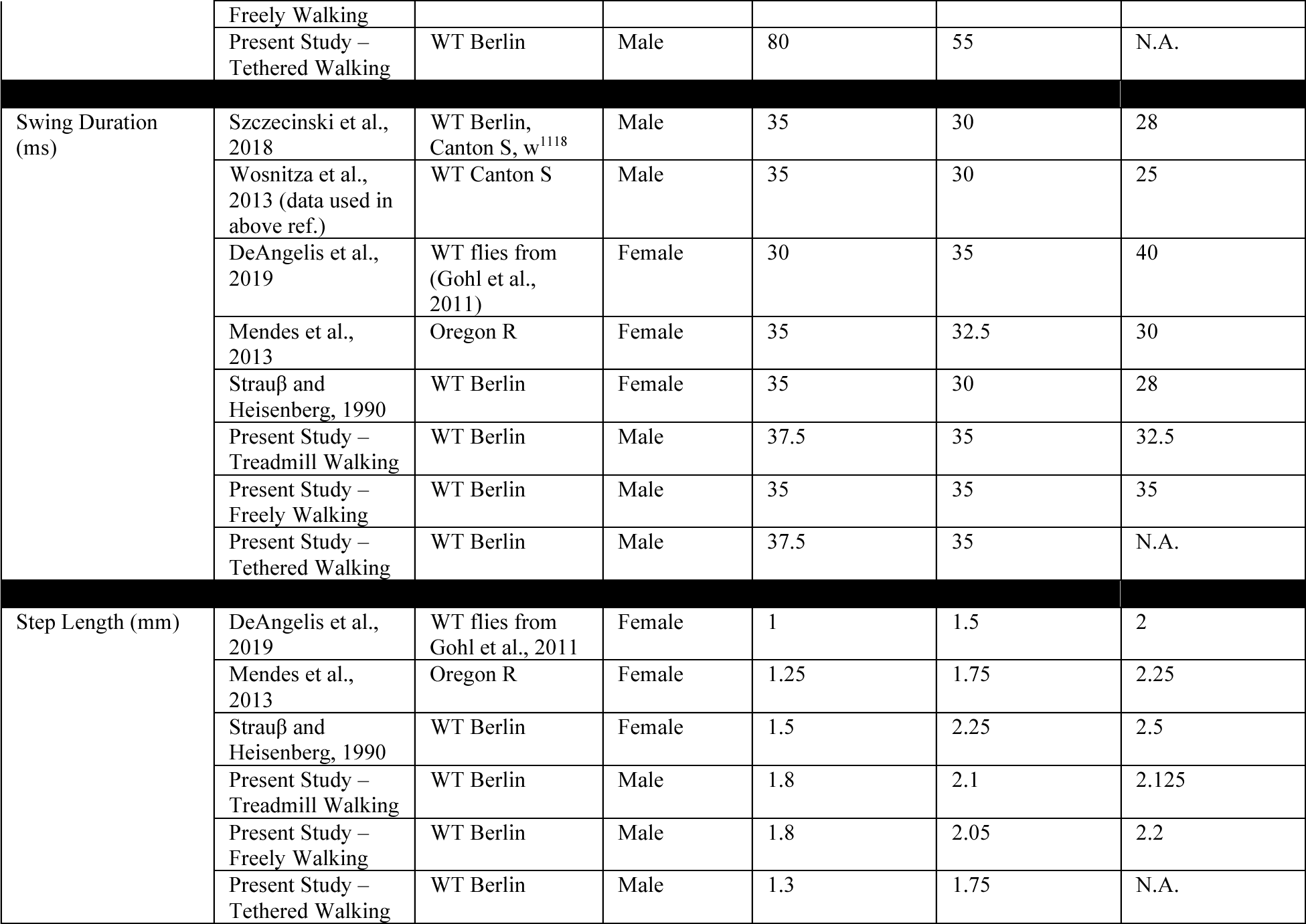
Summary of previously reported relationships between step kinematics and forward walking speed in *Drosophila melanogaster* and those reported in this study. Mean values were determined through visual inspection of fits within relevant plots of previous literature. The mean values for the relationships obtained in this study were approximated through visual inspection of the fits in Figure 2, as we did for the other papers.

## Video Legends

**Video 1. A representative fruit fly walking on the linear treadmill with a steady-state belt speed of ∼11 mm/s.** Top-down and side views of a wild-type Berlin fly walking on the treadmill within the chamber are shown by the top left and right videos, respectively. The videos were recorded at 180 fps. Below, in descending order, are the position of the fly within the chamber, its heading angle, its body velocity, and the belt speed throughout the video. The light blue regions indicates forward walking bouts. This data is displayed in **Figure 1D**.

**Video 2. Fruit flies are capable of walking across a wide range of treadmill belt speeds.** Top-down and side views of a wild-type Berlin fly walking on the linear treadmill within a chamber are shown by the top left and right videos, respectively. The tracked head (red), thorax (green), and abdomen (blue) are displayed for each camera view, as well as the belt speed profile contained within the video. Note that this early chamber design differs from that used in the rest of the paper.

**Video 3. The linear treadmill forces high-speed walking in flies.** Top-down and side views of a control (R52A01 DBD > tnt) fly walking on the linear treadmill within a chamber are shown by the top left and right videos, respectively. The fly was driven at a belt speed of about 18 mm/s and achieved walking velocities greater than 50 mm/s, which is the fastest walking velocity reported for *Drosophila melanogaster*. Note that flies of this genotype are not used elsewhere in the paper.

**Video 4. A fruit fly walking freely and forward in an arena.** The top-down view shows a wild-type Berlin fly walking in an arena (i.e. the fly bowl). The forward walking bout is indicated by the appearance of the “Forward Walking” label. The video was recorded at 150 fps and was slowed down 10x. This data is contained in **Figures 2 & 3**.

**Video 5. A representative tethered fruit fly walking on a sphere suspended by air.** Side and top-down views (left and right, respectively) show a wild-type Berlin fly walking on a floating sphere while tethered. The forward walking bout is indicated by the appearance of the “Forward Walking” label. The video was recorded at 300 fps and was slowed down by 5x. This data is contained in **Figures 2 & 3**.

**Video 6. Silencing chordotonal neurons alter walking in flies.** Top-down and side views (left and right, respectively) are shown for the same genetically-matched control fly (R52A01 DBD > Kir 2.1; top) and fly with chordotonal neurons broadly silenced (iav-GAL4 > Kir 2.1) as each walked on the linear treadmill. The belt speed was 17.9 mm/s for both flies. Videos were recorded at 180 fps and were slowed down by 2x. This data is contained in **Figure 4**

**Video 7. Flies use their middle legs to correct for rotational perturbations induced by asymmetric belt speeds on the split-belt treadmill.** Top-down views of the same control (R48A07 AD > Kir 2.1) fly walking on the split-belt treadmill when the left and right belts were tied in speed (10 mm/s) and when the left belt moved faster (12 mm/s) than the right (8 mm/s). The period of asymmetric belt movement is called the “split” period. During the representative split period, a forward walking bout is shown where the left and right legs of the fly walked on the corresponding belts. Videos were recorded at 200 fps and the video showing the forward walking bout was slowed 5x. This data is contained in **Figure 5**.

## Methods

### Fly Husbandry and Genotypes

Adult male *Drosophila melanogaster* between 2-7 days post-eclosion were used for experiments (Table 1). Flies were reared in a 25°C incubator with 14:10 light:dark cycle within vials filled with a standard cornmeal and molasses medium.

**Table 1.**
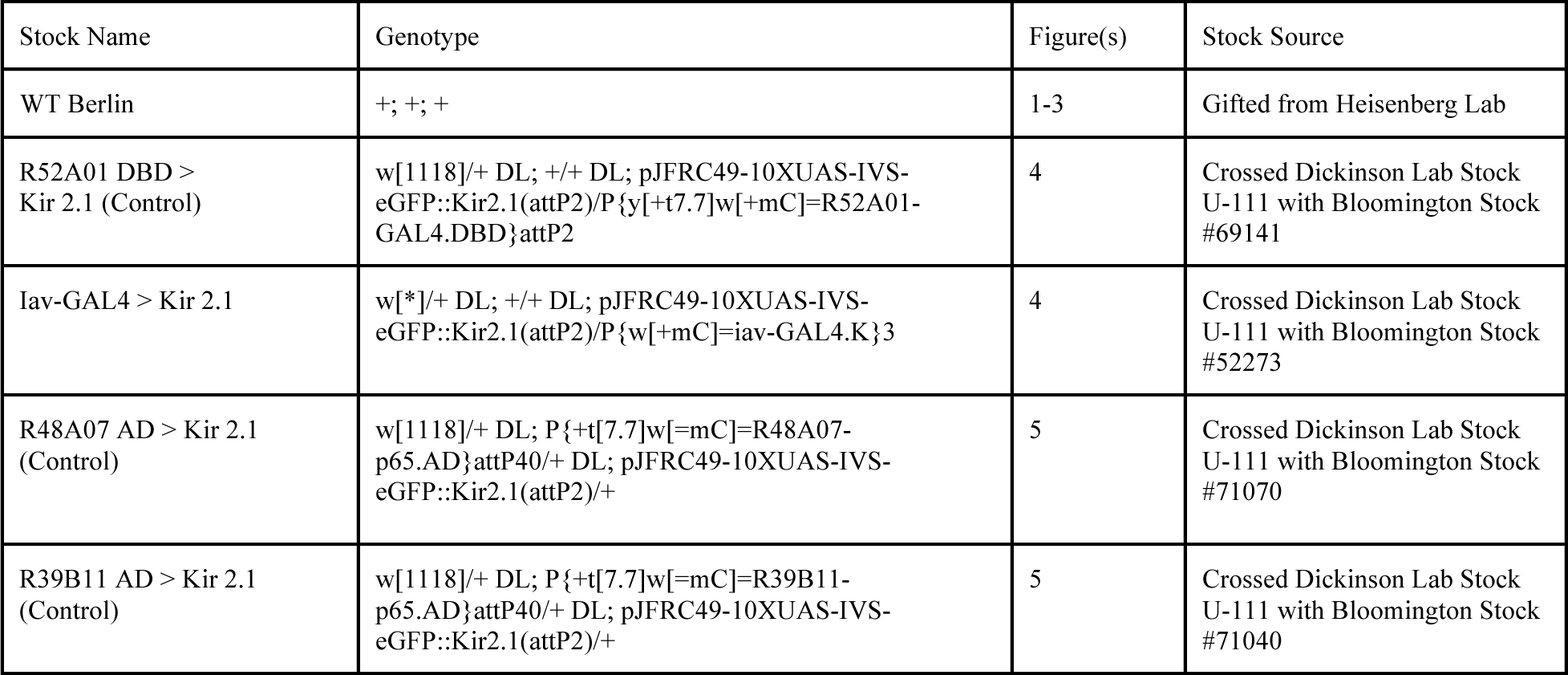
*Drosophila melanogaster* genotypes used for experiments.

### Miniature Treadmills

#### Linear treadmill and experiments

The key components of the linear treadmill system are a custom 3D-printed chamber, pulleys, a belt, a DC motor controlled programmatically with a PID controller, and 5 high-speed cameras. The chamber was designed using 3D CAD software (Autodesk Fusion 360: *Treadmill_Chamber.stl* in GitHub repository) and printed with black resin using a high spatial resolution 3D printer (Formlabs Form 2; black resin RS-F2-GPBK-04). The region of the chamber where a fly walked had transparent, sloped walls, a length of 8.929 mm, a max width of 6.5 mm, and a max height of 1.5 mm.

Coverslips that were 22m x 22m and size #1 were used as the transparent walls of the chamber (Electron Microscopy Sciences: #72200-10). Rain-X was applied to the inner surface of the coverslips to limit flies from walking on the glass. The pullies (B & B Manufacturing: 28MP025M6FA6) were attached to steel bars which rotated with bearings (AST Bearings: SMF126ZZ) that were mounted on custom fabricated brackets. The distance between the brackets was adjustable, which enabled the belt (B & B Manufacturing: 100MXL025UK) held between the pullies to be tensioned. A mounted DC motor (Phidgets: 12V/0.8Kg-cm/46RPM 50:1 DC Gear Motor w/ Encoder ID: 3256E_0) actuated the belt. An infrared ring-light (Olympus Controls: R130-850) was used to illuminate flies on the belt. 5 high-speed cameras (Machine Vision Store; USB 3.0 Basler camera acA800 x 600, Basler AG) with adjustable lenses (Computar: MLM3X-MP) and IR filters (Olympus Controls: FS03-BP850-34) recorded flies walking on the belt and within the chamber at 180 fps. The DC motor and high-speed cameras were controlled using a microcontroller (Phidgets: PhidgetMotorControl 1-Motor ID: 1065_1B) and DAQ (National Instruments: BNC-2110), respectively, and a custom Python script (*linear_treadmill_belt_stim_videography.py* in GitHub repository).

Male *Drosophila* (specific genotypes in Table 1) were driven to walk on the linear treadmill while the belt’s steady state speed was either 3.9, 7.4, 10.9, 14.4, or 17.9 mm/s. To smoothly reach steady-state belt speeds above 3.9mm/s, the belt linearly increased in speed with a slope of 3.5 mm/s^2^. Wild-type Berlin and iav-GAL4 > Kir 2.1 flies were subjected to each belt speed 10 times, whereas R52A01 DBD > Kir 2.1 flies were presented each belt speed 15 times. Trials in which flies walked at a given belt speed were 10 seconds. There was a 5 second period between trials where the belt moved at 3.9 mm/s and the high-speed videos were saved. DeepLabCut and Anipose (Karashchuk et al., 2021; Mathis et al., 2018) were used track the fly’s leg tips (i.e. tarsi), head, thorax, abdomen, and key points on the chamber in 3D using 2,250 annotated frames as the training dataset. The test prediction error of the tracking was 5.45 pixels and the reprojection error was 2.88 pixels. Walking kinematics were analyzed and visualized using custom Python scripts (*linear_treadmill_walking_analysis.ipynb* & *linear_treadmill_visualization_walking_comparisons.ipynb* in GitHub repository).

We also tried driving tethered flies to walk on the linear treadmill. Occasionally, the front legs moved with the belt, but the overall movement between legs was uncoordinated. We typically observed legs being dragged along the surface of the belt. It should be noted that we tried many different tether designs, from rigid ones to light-weight, low-resistance ones inspired from a treadmill used for desert ants (Dahmen et al., 2017). Overall, we were unable to drive coordinated walking in tethered flies using the treadmill.

#### Split-belt treadmill and experiments

The construction of the split-belt chamber was similar to the linear treadmill. The key difference was the addition of a second independently actuated belt. Therefore, we used smaller belts (B & B Manufacturing: 100MXL012UK) and pullies (B & B Manufacturing: 28MP012M6FA6), while the chamber size remained the same. We also used DC motors (Phidgets: 12V/3.0Kg-cm/78RPM 51:1 DC Gear Motor w/ Encoder ID: 3263E_1) that were of a newer model. Finally, the frame rate of the high-speed cameras was increased to 200 fps. A custom Python script controlled the motors and cameras (*splitbelt_treadmill_belt_stim_videography.py* in GitHub repository).

Only male flies were used in split-belt experiments (specific genotype in Table S1). Flies initially walked on belts that were tied in speed (i.e. 10 mm/s) for 10 minutes. Then, one belt increased in speed by 20% (i.e. 12 mm/s) while the other belt decreased in speed by 20% (i.e. 8 mm/s). This split period also lasted 10 minutes. Following the split period, the belts again moved at 10 mm/s for 10 minutes. At the end of the 10 minutes, this trial structure was repeated, but the belts switched which increased or decreased in speed during the split period. 5 high-speed cameras recorded the movement of the fly during this task and the same key points in the linear treadmill experiments were tracked and reconstructed in 3D.

The training dataset consisted of 4,140 annotated frames and the DeepLabCut network achieved a test error of 6.13 pixels. The reprojection error was 1.82 pixels. Custom Python scripts were used to analyze and visualize walking kinematics (*splitbelt_walking_analysis.ipynb* and *splitbelt_walking_visualization.ipynb* in GitHub repository).

### Freely Walking Experiments

Groups of 10, 2-7 day old, male wild-type Berlin flies were placed in a 10 cm circular arena with sloped walls (i.e. fly bowl) and allowed to freely walk (Simon and Dickinson, 2010). A high-speed camera (Machine Vision Store: USB 3.0 Basler camera acA1300-200um, Basler AG) recorded a 2.4 cm x 2.7 cm region of the arena from above at 150 fps in 10s bouts. Leg tips, head, thorax, and abdomen of flies were tracked using SLEAP, which is optimized for multi-animal pose estimation (Pereira et al., 2022). Custom Python scripts quantified and visualized walking kinematics (*freely_walking_analysis_visualization.ipynb* in GitHub repository).

### Tethered Experiments

De-winged male wild-type Berlin flies, 2-5 days old, were attached to a thin tungsten tether (0.1 mm) with UV curing glue (KOA 300). Flies were then positioned with a micromanipulator on a spherical foam ball (weight: 0.13 g; diameter: 9.08 mm) suspended by a regulated air supply. The 2D trajectory, and forward, rotational, and side-slip velocities of the fly were measured from the movement of the ball with FicTrac (Moore et al., 2014). 6 high-speed cameras (Machine Vision Store: USB 3.0 Basler camera acA800 x 600, Basler AG) recorded flies walking on the ball at 300 fps over 2 second bouts. Custom python and MATLAB scripts were used to acquire the high-speed videos. The leg joints, tips, head, and abdomen were tracked and reconstructed in 3D using DeepLabCut and Anipose, respectively. Walking kinematics were analyzed and visualized using Python (*tethered_walking_analysis_visualization.ipynb* in GitHub repository).

### Statistical and KL Divergence Analyses

Chi-squared test and t-tests were used to test for differences in step kinematics between the different experimental setups (**Figure 2**). Statistics were conducted on kinematic distributions containing data from all leg pairs and that were associated with a walking speed between 6.5-10.1 BL/s. This walking speed range was chosen because it contained 50% of the data across setups given their overlapping speed ranges. The chi-squared test determined whether the proportion of values of a given discretely measured step metric (i.e. step frequency, stance duration, and swing duration) was the same between freely walking flies and those in the other two setups. A Bonferroni correction of 18 was added to account for multiple comparisons (6 legs and 3 setups). Therefore, a significant difference was determined to be p < 0.0028. Note that to make the statistical comparisons of step frequency, stance duration, and swing duration between all three setups, we had to interpolate the underlying signal for treadmill and freely walking flies from 180 fps and 150 fps, respectively, to 300 fps (i.e. the tethered setup sampling rate). Finally, a t-test was used to determine significant differences between the mean step lengths of freely, treadmill, and tethered walking flies. A Bonferroni correction of 18 was also applied.

To determine whether step kinematics of freely walking flies were more similar to those of treadmill or tethered walking flies, we computed the relative KL divergence (**Figure S1**). KL divergence computes an unbounded similarity, in the form of entropy, between two distributions, where a value closer to zero indicates greater similarity. Thus, we computed the KL divergence between freely and treadmill step kinematic distributions, and freely and tethered distributions. Then, we calculated a relative similarity score between the two sets of distributions by using the following equation:

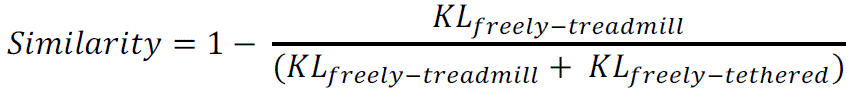

A value of 1 indicated that freely and treadmill walking step kinematics were more similar than freely and tethered ones, and vice versa.

Finally, a bootstrap statistical analysis was used to test if there were significant shifts in the mean anterior and posterior extreme positions across the different belt conditions (i.e. tied, slow, and fast belt) of the split-belt task. A Bonferroni correction of 36 was applied (6 legs, 3 belt conditions, and 2 axes of comparison), requiring a p < 0.0014 for a significant difference.

### Kinematic Classification and Parameters

#### Swing and stance classification

Leg tip velocity was used to classify leg swing and stance phases in all walking setups. We first computed the instantaneous speed of each leg tip from their allocentric positions along the longitudinal and lateral body axes. For the treadmill and tethered walking setups, a rotation matrix had to be applied to the position data to align all flies to a common reference frame and to ensure symmetric contralateral leg movement with respect to the defined axes. The instantaneous speed was then transformed into velocity by applying a negative sign to instances where the leg moved backward along the longitudinal body axis in an egocentric reference frame. This forced the stance to be negative. To achieve this sign application for freely and treadmill walking setups, where the heading of the fly is constantly changing, we rotated the fly in each frame to a common heading. This made it easier to distinguish the period when the leg moved along the body. The instantaneous velocities of the leg tips were then smoothed with a Gaussian kernel. The width of the kernel was chosen such that the signal wasn’t oversmoothed, but instantaneous tracking errors were mitigated. Swing was classified as periods where the smoothed leg tip velocities were above and below manually chosen upper (treadmill: 5 mm/s, freely: 15 mm/s, tethered: 0 mm/s) and lower thresholds (all setups: –25 mm/s). Stance was classified as the period where the leg tip velocities were between these thresholds. Finally, we corrected blips in the classification (e.g. converting a swing period consisting of 1 frame into stance) and matched the stance and swing onsets of a given step.

We checked the accuracy of the swing and stance classifications by manually inspecting the raw high-speed videos. Note that we also tried to perform swing and stance classification by thresholding the Hilbert transformed longitudinal body axis position signal of each leg tip, and doing peak detection on that signal, but both methods performed more poorly than the method described above. The Hilbert transform assumes that a signal is non-stationary, which is invalid when using a leg position signal that dynamically moves in 3D. Peak detection also fails to compensate for the richness of leg movement. Overall, our accurate classifications of swing and stance enabled precise quantifications of step kinematics and inter-leg coordination.

#### Forward walking bout classification

Different forward walking bout classifiers were used for each walking setup. In the linear and split-belt setups, forward walking bouts were periods lasting 200 ms where the fly walked in the middle of the chamber, had a heading angle within –15 to 15 degrees with respect to the front of the chamber, and had a forward walking velocity (aligned to the driving axis of the treadmill) greater than 5 mm/s. Flies were classified as walking in the middle of the chamber if their abdomen was 1.85 mm in front of the back of the chamber and the tarsi of their front legs were 1.08 mm behind the front of the chamber. For freely walking flies, the thorax position, specifically the angle between sets of 3 position sample points, was used to first isolate straight body trajectories in allocentric coordinates. A straight trajectory was defined as one where the angles between thorax position sample points were less than 4.5 degrees. Once a straight trajectory was isolated, we classified it as a forward walking bout if the corresponding fly’s average body velocity was greater than 5 mm/s, the inter-quartile range of the heading angles was less than 20 degrees, and the duration of the trajectory was greater than 200 ms. Lastly, forward walking bouts of tethered flies were first identified by using a previously described behavioral classifier (Karashchuk et al., 2021), but later refined to those greater than 200 ms in duration, having an average forward velocity greater than 5 mm/s, a minimum instantaneous forward velocity of 0.5 mm/s, an average absolute rotational velocity less than 25 degrees/s, and an absolute instantaneous rotational velocity less than 100 degrees/s. Across all setups, the first and last steps of all legs were trimmed within identified forward walking bouts to compensate for the transitions into and out of them.

#### Forward walking step filtering

Forward walking steps were filtered based on step frequency, stance duration, and swing duration in all walking setups. Forward walking steps were considered to be those that had a step frequency between 5 and 20 steps/s, a swing duration between 15 and 75 ms, and a stance duration less than 200 ms. These filtering thresholds were empirically determined and based on previously published results of forward walking step kinematics in fruit flies (DeAngelis et al., 2019; Mendes et al., 2014; Szczecinski et al., 2018; Wosnitza et al., 2013).

## Glossary of kinematic parameters

Body length: the distance between the head and distal part of the abdomen.
Body height: the vertical distance between the ground and thorax.
Step frequency: the number of steps completed within a second.
Stance duration: the duration that a leg contacts the ground while walking.
Swing duration: the duration of the aerial phase of leg movement during walking.
Step length: the total distance a leg travels within a step (i.e. stance onset to the subsequent stance onset) in allocentric coordinates.
Step distance: the total distance a leg travels within a step (i.e. stance onset to the subsequent stance onset) in egocentric coordinates.
Step speed: the total distance a leg travels within a step in egocentric coordinates divided by the duration of the step.
Anterior extreme position: the position where a leg first contacts the ground (i.e. stance onset) in egocentric coordinates.
Posterior extreme position: the position where a leg first takes off from the ground (i.e. swing onset) in egocentric coordinates.
Number of legs in stance: the number of legs contacting the ground at a given moment in time.
L1 relative phase: the relative offset in the stance onsets between the left front leg and the leg of interest with respect to the left front leg’s step cycle.
Tripod step order: the order in which the legs within a tripod group (i.e. ipsilateral front and hind legs and the contralateral middle leg) first enter stance with respect to the left front leg’s step cycle.

